# PiDeeL: Pathway-Informed Deep Learning Model for Survival Analysis and Pathological Classification of Gliomas

**DOI:** 10.1101/2022.10.21.513161

**Authors:** Gun Kaynar, Doruk Cakmakci, Caroline Bund, Julien Todeschi, Izzie Jacques Namer, A. Ercument Cicek

**Affiliations:** Computer Engineering Department, Bilkent University, 06800, Ankara, Turkey; School of Computer Science, McGill University, Montreal QC, Canada; MNMS Platform, University Hospitals of Strasbourg, France; ICube, University of Strasbourg, CNRS UMR, 7357, Strasbourg, France; Department of Nuclear Medicine and Molecular Imaging, ICANS, Strasbourg, France; Department of Neurosurgery, University Hospitals of Strasbourg, Strasbourg, France; Computational Biology Department, Carnegie Mellon University, PA, 15213, Pittsburgh

**Keywords:** Glioma, Metabolomics, NMR, Survival Analysis, Intraoperative Feedback, Deep Learning

## Abstract

Online assessment of tumor characteristics during surgery is important and has the potential to establish an intraoperative surgeon feedback mechanism. With the availability of such feedback, surgeons could decide to be more liberal or conservative regarding the resection of the tumor. While there are methods to perform metabolomics-based online tumor pathology prediction, their model complexity and, in turn, the predictive performance is limited by the small dataset sizes. Furthermore, the information conveyed by the feedback provided on the tumor tissue could be improved both in terms of content and accuracy. In this study, we propose a metabolic pathway-informed deep learning model, PiDeeL, to perform survival analysis and pathology assessment based on metabolite concentrations. We show that incorporating pathway information into the model architecture substantially reduces parameter complexity and achieves better survival analysis and pathological classification performance. With these design decisions, we show that PiDeeL improves tumor pathology prediction performance of the state-of-the-art in terms of the Area Under the ROC Curve (AUC-ROC) by 3.38% and the Area Under the Precision-Recall Curve (AUC-PR) by 4.06%. Similarly, with respect to the time-dependent concordance index (c-index), we observe that PiDeeL achieves better survival analysis performance (improvement up to 4.3%) when compared to the state-of-the-art. Moreover, we show that importance analyses performed on input metabolite features as well as pathway-specific hidden-layer neurons of PiDeeL provide insights into tumor metabolism. We foresee that the use of this model in the surgery room will help surgeons adjust the surgery plan on the fly and will result in better prognosis estimates tailored to surgical procedures.

**Availability:** The code is released at https://github.com/ciceklab/PiDeeL. The data used in this study is released at https://zenodo.org/record/7228791.

**Supplementary information:** Supplementary data are available at *Briefings in Bioinformatics* online.

## Introduction

Gliomas have uncertain patient mortality rates due to differences in suspected brain cell types and the variance observed in cancer progression dynamics [38]. 2021 version of central nervous system tumor taxonomy developed by the World Health Organization (WHO) [40] categorizes gliomas according to cell types (e.g. astrocytes, oligodendrocytes) as well as four pathologically different grades (i.e. grade I - IV). This taxonomy takes differential histopathologic, molecular, and genetic characteristics of gliomas into account and orders grades with respect to increasing malignancy and decreasing prognostic optimism. Considering the prevalence of gliomas among brain tumors diagnosed in adults [41], devising a surgical management strategy that aims to make personalized clinical decision-making plausible via an intraoperative or preoperative surgeon feedback mechanism becomes essential.

In order to amplify the percentage of tumor tissue resected from a patient’s brain, various imaging [49], spectroscopy as well as mass spectrometry [4, 46, 15, 42, 45, 9, 10, 26, 19], and optical spectrometry [48, 51, 12, 36, 33, 34, 28, 27, 11, 39, 44, 24, 54] based techniques, were proposed in the literature. As a member of this cohort, ^1^ H High-Resolution Magic Angle Spinning Nuclear Magnetic Resonance (HRMAS NMR) spectroscopy measures metabolic activity in a given unprocessed tissue specimen [22] with a turnaround time of *∼* 20 mins and generates a time domain signal (Free Induction Decay - FID) in the form of the sum of decaying exponential functions. The FID signal is often preprocessed to obtain the HRMAS NMR spectrum defined on the frequency domain.

After preprocessing, the HRMAS NMR spectrum becomes well-suited to analyze the presence of left-over tumor tissues in the excision cavity and provide feedback to the surgeon about the nature of the tumor. Establishing this HRMAS NMR-guided intraoperative feedback loop requires a pathologist knowledgeable about tumor metabolism, and an NMR technician who detects an abundance of a few metabolites from the spectrum. However, the feedback resulting from this strategy is limited by (1) the availability of the experts during the surgery timeframe; and (2) the throughput of the metabolite quantification procedure manually performed by the technician one metabolite at a time for HRMAS NMR spectra obtained from each tissue specimen. Moreover, overlapping metabolite peaks may not be fully separated in the preprocessed ^1^ H spectra [29].

Due to the high-dimensional, complex, and structured nature of the preprocessed HRMAS NMR spectra, data-driven computational modeling (i.e. machine learning) has the promise to be a valuable companion to the above-mentioned manual feedback loop. To this end, various supervised machine learning models were explored for providing automated feedback on pathological classification [7, 8] of the resected glioma tissue specimens. The dataset used for these studies [6] consists of HRMAS NMR spectra of 568 glioma and healthy control tissue samples. It is relatively small despite being the largest of its kind which prohibited the use of complex models and forced the models built to be shallow. This suggests room for improvement in the performance of the feedback pipeline.

From a methodological perspective, the applicability of deep learning models in small data regimes can be improved by making informed architectural choices to simplify models. Incorporating biological prior information may limit the connectivity of the neurons in a model and simultaneously (1) alleviate overfitting when available dataset sizes are small, and (2) aid model interpretability regardless of the dataset size. While there is no application in the metabolomics domain, biologically informed deep learning architectures were successfully utilized in the context of metastasis prediction in patients with prostate cancer based on their genomic profiles [17], and phenotype prediction based on the genetic variants of the patients [52].

Furthermore, the pathological classification of tumors may not be the only determinant of multi-faceted and patient-specific cancer progression semantics. In the case of glioblastoma multiforme (GBM), a grade IV astrocytic glioma type, a dismal prognosis is often expected due to the highly malignant and invasive nature of this class of tumors [38]. Despite their severity, prolonged and progression-free survival was observed in isocitrate dehydrogenase (IDH) wildtype GBM [43, 5, 53]. This finding promotes the inclusion of survival risk estimates in the feedback given to surgeons for more accurate surgical decision-making.

Here, we propose a deep neural network architecture (PiDeeL) to predict the survival of the glioma patient and the pathological classification of the tumor using the HRMAS NMR signal to give feedback to the surgeons (Figure 1). While there were methods proposed for automated pathological classification, this is the first study to incorporate these two orthogonal sources of information to the surgeon as feedback at the same time. For the first time in the metabolomics domain, we incorporate prior biological information from the metabolic networks into the neural network architecture and increase the interpretability of the predictions which is also important for the surgeon to understand the underlying biology. We show that PiDeeL achieves better performance compared to the state-of-the-art model in pathological classification and, provides survival prediction with a c-index of 68.7%. We also analyze the most important metabolites and metabolic pathways for the reasoning of the model and discuss their relevance in the context of gliomas.

**Fig. 1.**
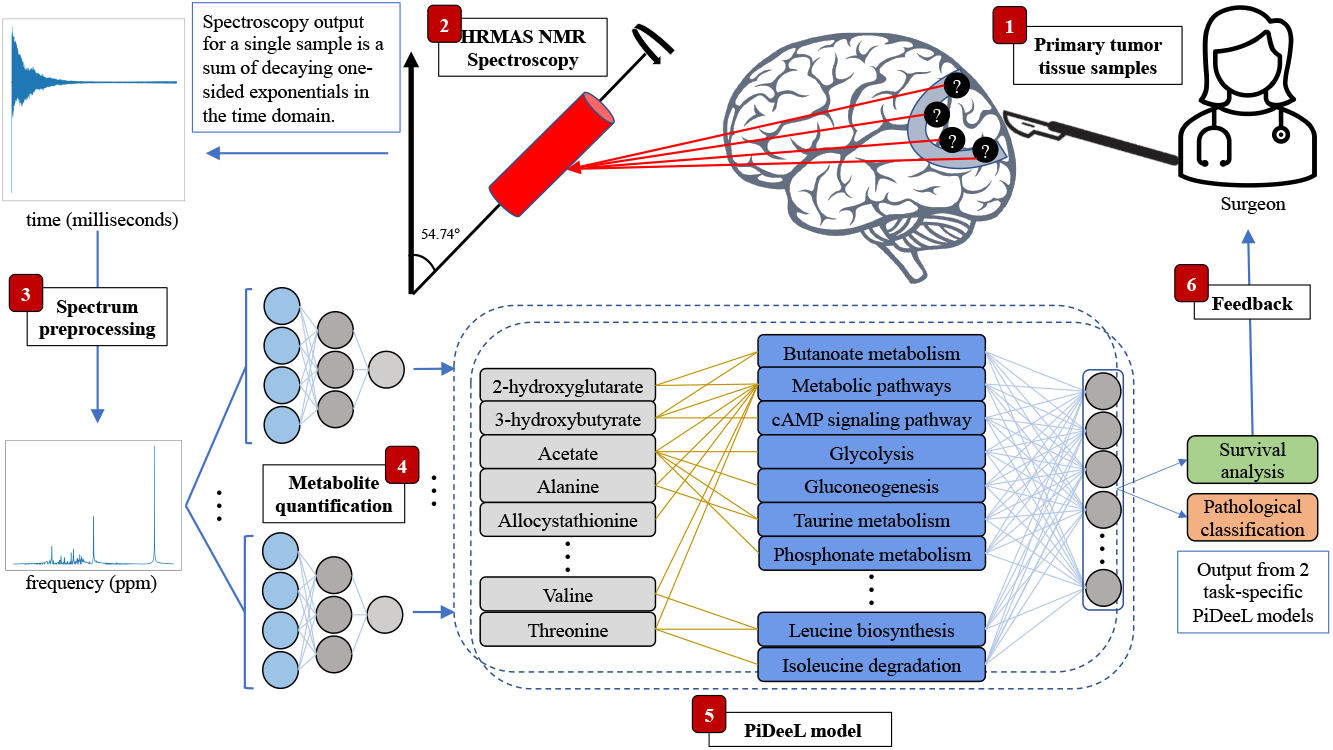
PiDeeL assisted intraoperative feedback loop. (Step 1) The surgeon removes primary tumor tissue from the patient’s brain and prepares specimens from the resected tissue. (Step 2) Prepared samples are sent for HRMAS NMR spectroscopy analysis. (Step 3) Time domain signals resulting from the spectroscopy of each sample are preprocessed to uncover metabolite peaks harbored in the corresponding frequency domain spectrum. (Step 4) 37 metabolites are quantified from preprocessed spectra in the order of milliseconds via metabolite-specific 2-layer fully-connected neural networks. (Step 5) PiDeeL transforms metabolite concentrations to survival analysis and pathological classification predictions also in the order of milliseconds. (Step 6) Based on this feedback, the surgeon decides to preempt or continue performing the surgery, again with the guidance of PiDeeL.

## Methods

### Dataset

The dataset used in this study is a subset of the patient cohort released at https://zenodo.org/record/5781769, and it consists of 568 HRMAS NMR spectra acquired from 458 glioma (80.7% aggressive and 19.3% benign) and 110 control tissue specimens collected during respective surgeries of 513 patients performed at University Hospitals of Strasbourg. For the details of tissue specimen collection, and ethics statement, please see Section 1.1 of the Supplementary Text.

In order to prepare the dataset for survival analysis, we retain the samples acquired from primary tumor tissue specimens of glioma origin whose record of the date of metabolomics-guided surgery and the date of decease or last control is present. We removed specimens resected from healthy control patients and excision cavities of glioma patients. The resulting set of 384 HRMAS NMR spectra contains one sample per glioma patient and is composed of 299 (77.9%) aggressive and 85 (22.1%) benign tumors whose subtype and grade information are provided in Supplementary Table 1. 70.1% of the patient cohort (269 patients) was reported deceased before the spectra acquisition from tissue specimens (i.e. until January 2021). The remaining 115 patients were considered to be censored as their survival status is not known. The distribution of patient age and time-to-event for deceased and censored patients are provided in panels A, B, and C of Supplementary Figure S1, respectively. Furthermore, ERETIC- CPMG HRMAS NMR FID signals of these 384 glioma patients, whose spectra were acquired according to the procedure described in Section 1.1.3 of Supplementary Text, are released at https://zenodo.org/record/7228791.

### Automated Metabolite Quantification

In this section, we describe the steps applied to the FID signals (output of HRMAS NMR Spectroscopy) to obtain metabolite concentrations, which were used as input to the survival analysis and pathological classification models presented throughout the work. We adopt the deep learning-based automated metabolite quantification method proposed in Cakmakci et al. (2022) in order to quantify 37 metabolites of interest from HRMAS NMR spectra. Briefly, we preprocess our samples according to the input specification of their pipeline and quantify each metabolite from the spectra in our sample set with their publicly available quantification models per metabolite from https://github.com/gunkaynar/targeted_brain_tumor_margin_assessment. For the complete list of 37 metabolites, details of automated metabolite quantification, and HRMAS NMR spectrum preprocessing, please see Section 1.2 of Supplementary Text.

Formally, let the predicted concentration of a given metabolite *j* for a sample set of size *N* be 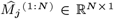. We denote the full output of the automated metabolite quantification pipeline for 37 metabolites and a sample set of size *N* as the concatenation of all predicted quantification vectors across metabolites:

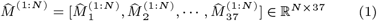

### Problem Formulation

Here, we formulate pathological classification and survival analysis tasks based on the full output of the metabolite quantification pipeline described in the Automated Metabolite Quantification Section as well as tumor malignancy label, patient-specific survival event indicator (i.e. censored or deceased), and time-to-event variables. For a given sample *i*, we denote the feature vector, tumor malignancy label, survival event indicator, and time-to-event as 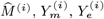 and 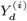, respectively. The feature vector, 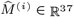, is the 37-dimensional vector corresponding to the full output of the automated metabolite quantification pipeline. Survival event indicator, 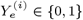, is a binary variable that indicates whether the sample is from a deceased (i.e. 1) or a censored patient (i.e. 0). Time to survival event, 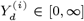 is a non-negative continuous variable defined as the time (in days) between metabolomics-guided tumor removal surgery and the date of decease or last control for deceased and censored patients, respectively. Finally, the tumor malignancy label, 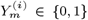, indicates whether the sample is from a tumor having aggressive (i.e. 1) or benign (i.e. 0) pathology, similar to the problem formulations presented in Cakmakci et al. (2020, 2022).

For our formulation of the survival analysis task, we refer to the hazard function denoted as 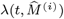 for a given sample *i* where *t* is time-to-event (i.e. 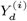). Assuming proportional hazards from Cox’s model [13], the hazard function factorizes [18, 30] to a time-dependent baseline hazard function, *h*_0_ (*t*), and a sample-dependent risk function, 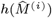, as given in Equation 2.

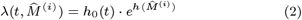

We train survival analysis models based on this formulation of hazard function on a dataset of *N* samples to learn a mapping function *f* : ℝ^37^ → [*−∞, ∞*] such that 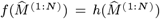. Here, the goal of the trained models is to predict risk scores for glioma patients based on their metabolic profile. In order to infer these models, we make use of 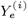 and 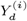 as labels.

On the other hand, we train pathological classification models on a dataset of *N* samples to learn a mapping function *g* : ℝ^37^→ 0, 1 such that 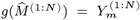 (a binary classification problem similar to Cakmakci et al. (2020, 2022).

### Baseline Models

In this section, we provide the details of models benchmarked against PiDeeL for survival analysis and pathological classification tasks formulated in the Problem Formulation Section.

#### Survival Analysis

For the survival analysis task, we consider Cox Proportional Hazards (Cox-PH), Random Survival Forest (RSF), and DeepSurv [30] models as baselines.

Cox-PH model is a linear model that assumes proportional hazards to factorize instantaneous hazard function as a product of a baseline hazard function and a sample-dependent risk function, 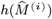, as formulated in the Problem Formulation Section. Based on the formulation provided by Cox et al. (1972), the risk function estimate for sample *i*, 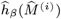, is provided by the linear model given in Equation 3.

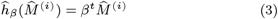

where 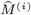 is the feature vector of metabolite concentrations for sample *i*, and *β* is the coefficient vector. The model is fit by maximizing Cox partial likelihood [18, 30, 13] given in Equation 4.

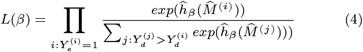

RSF [25] is a nonlinear ensemble method that constructs a predefined number of survival trees using a random subset of features and bootstraps of training data to estimate the cumulative hazard function, Λ(*t*), defined in Equation 5. Internal nodes of survival trees are split in order to maximize log-rank test [47, 32] statistic between resulting child nodes.

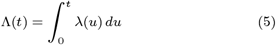

Neural network-based approaches to survival analysis are introduced by Faraggi et al. (1995). Deeper variants of these models are more recently proposed as DeepSurv model [30] which is a multilayer perceptron that models sample-specific risk function, 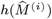, as formulated in the Problem Formulation Section. The model has a single output neuron with linear activation and adopts a negative log of Cox partial likelihood given in Equation 6 as the loss function.

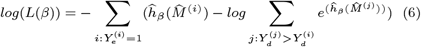

We consider 2, 3, and 4-layer DeepSurv as our baseline models. We use a single neuron in the last layer with linear activation to output the predicted risk scores of samples. The trained 3-layer DeepSurv model on our dataset of size *N* can be summarized as follows:

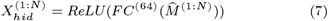

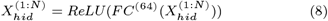

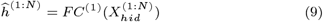

where *X*_*hid*_ represents the output of the first hidden layer, 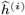 represents the predicted risk score, *FC*^(*·*)^ represents a fullyconnected layer with (*·*) neurons, and *ReLU* represents the rectified linear unit. In the case of 2 and 4-layer DeepSurv models, we remove or repeat the hidden layer defined in Equation 8.

#### Pathological Classification

We consider random forest (RF) from Cakmakci et al. (2022) and fully-connected neural networks as baseline models for the pathological classification task.

We consider neural network-based models for the pathological classification task as baseline models. We train 2, 3, and 4-layer fullyconnected neural networks with class-weighted binary cross entropy loss. The trained 3-layer fully-connected networks on our dataset of size *N* can be summarized as follows:

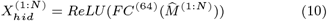

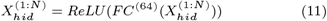

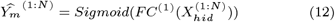

where *X*_*hid*_ represents the output of the first hidden layer, 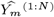 represents the predicted tumor pathology, *FC*^(*·*)^ represents a fullyconnected layer with (*·*) neurons, and *ReLU* represents the rectified linear unit. We also consider 2 and 4-layer fully-connected networks where we remove or repeat the hidden layer defined in Equation 11, respectively.

Pathway-Informed Deep Learning Model (PiDeeL)

In this section, we introduce PiDeeL, a pathway-informed multilayer perceptron model designed to integrate biomarker metabolite concentrations provided as input at the metabolic pathway level. We use the Kyoto Encyclopedia of Genes and Genomes (KEGG) database to parse the metabolite-to-pathway mapping and construct a 37×138 matrix, hereafter *PI matrix*, where the value in the *i*^*th*^ row and *j*^*th*^ column of matrix *PI matrix* is 1 if the metabolite *i* appears in the pathway *j*, and 0 otherwise. Similar to the baseline fully-connected neural network models for the survival analysis and pathological classification tasks, we consider 2, 3, and 4-layer PiDeeL models.

#### Model Architecture

Given a dataset of *N* samples, the architecture of the 3-layer PiDeeL model for the survival analysis can be summarized as follows:

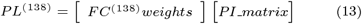

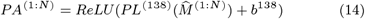

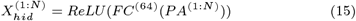

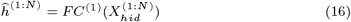

where *PL*^(138)^ and *b*^138^ represent the weights and biases of the pathway-informed layer, *PA*^(1:*N*)^ represents the activations of the first hidden layer (hereafter pathway-activation), *X*_*hid*_ represents the output of hidden layers, 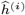 represents the predicted risk score, *FC*^(*·*)^ represents a fully-connected layer with (*·*) neurons and *ReLU* represents the rectified linear unit. We incorporate the metabolic pathway information into PiDeeL by defining the first layer of the model using *PI_matrix*. This architectural decision makes the first layer of PiDeeL sparse since the connections in the first layer are only permitted to be formed between input metabolites and the pathways they are involved in. We use the negative log of Cox Partial Likelihood as the loss function, similar to the DeepSurv models formulated in Survival Analysis Section, in order to train PiDeeL for the survival analysis task.

The architecture of PiDeeL used for survival analysis and pathological classification tasks differs only in the output layer (i.e. Equation 16). In order to adapt PiDeeL for pathological classification we use the sigmoid function as the output activation function by replacing Equation 16 with Equation 17. Additionally, we use class-weighted binary cross entropy as the loss function.

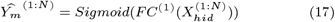

#### Interpretability and Importance Analyses

We perform SHapley Additive exPlanations (SHAP) analysis on our feature vector 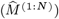, and pathway activations (*PA*^(1:*N*)^) to assess the importance of each metabolite and pathway on the prediction of PiDeeL. For the details of these analyses, please see Section 1.4.3 of the Supplementary Text.

#### Ablation Studies for Pathway-Informed Architecture

In order to verify that the improvements achieved by the PiDeeL model over baseline models are due to the pathway-informed architecture, we conduct a series of ablation tests as follows:

- We train fully-connected models with 138 neurons in the first hidden layer to check whether the number of neurons in the first layer (138) of PiDeeL contributes to the performance.
- We train PiDeeL models using randomly connected *PI_matrix* instead of *PI matrix* that are constructed from the KEGG database to check whether the pathway-informed connections have a benefit over the same number of randomly-connected edges.
- We train PiDeeL models using shuffled *PI_matrix*. This is similar to randomly connected *PI_matrix* except it preserves the in-degrees of pathways.
- We train the DeepSurv model with different dropout rates (0.5, 0.6, 0.7, 0.8, 0.9) in the first hidden layer to check whether the pathway-informed layer has a benefit over this widely used technique to regularize neural networks.

Please refer to Section 1.4.4 of the Supplementary Text for the details of ablation studies for pathway-informed architecture based on survival analysis.

## Results

### Experimental Setup

In this section, we describe the experimental setup used for evaluating PiDeeL as well as baseline models formulated in the Baseline Models Section in the context of the survival analysis scenario. Please see Section 1.4.1 of the Supplementary Text for the results of the pathological classification. We measure the performance of survival analysis models in the unit of time-dependent concordance index (c-index) described in Antolini et al. (2005). The original concordance index [23] is computed with the individually predicted patient risk scores, whereas, with a time-dependent definition, individual risk scores are calculated with the predicted risk function and the baseline hazard function that is computed on the sample set [3, 21].

For each task and model type, we adopt a 5-fold cross-validation routine and repeat it 3 times with different seeds (i.e. 15 iterations). Before each iteration, we shuffle and divide our dataset into training, validation, and test set such that no sample or patient is shared across folds. The results discussed hereafter are with respect to the predictive capability of models on the test split. During iteration *i* of the cross-validation setup, Cox-PH and RSF models were tested on the fold *i*, validated on fold (*i*+1)*mod*5 and trained on the remaining 3 folds. We re-train the selected configuration on the union of training and validation splits to get stronger baseline models. For all deep learning models, re-training steps are not performed due to the presence of early-stopping dynamics. Please see Section 1.3 of Supplementary Text for hyperparameter space and selection details.

### Survival Analysis

Please see Figure 2 for the survival analysis performance of CoxPH and RSF (gray boxplots) as well as DeepSurv (cyan boxplots) and PiDeeL (navy blue boxplots) with respect to the c-index. We observe that 4-layer PiDeeL outperforms baselines and DeepSurv models on survival prediction and achieves a median c-index of 68.7%. We also see that the performance slightly increases when the depth of PiDeeL is increased from 2 to 4 (improvement up to median c-index of 0.4%). On the other hand, the performance of DeepSurv models diminishes with increasing model depth (deterioration up to a median c-index of 1.5%). Furthermore, when DeepSurv and PiDeeL with the same number of layers are compared, we see the performance gap increase with respect to the number of layers in the model. Here, we quantitatively observe the benefit of incorporating biological information in modeling choices. For 2, 3, and 4-layered networks, PiDeeL improves upon DeepSurv with the same model depth by 2.5%, 3.4%, and 4.3% according to the median c-index, respectively. This supports the robustness of sparse architectures in small data regimes for the survival analysis task. When the performance of baseline models is considered in this setup, RSF appears to be a particularly strong baseline with a median c-index of 67.8%. This is also expected since the power of random forest-based models on HRMAS-NMR data has been demonstrated previously on a cohort of similar size [7, 8]. Still, PiDeeL achieves better performance than RSF also providing more interpretability by enabling pathway-activation analysis. Finally, the Cox-PH model performs substantially worse than PiDeeL.

**Fig. 2.**
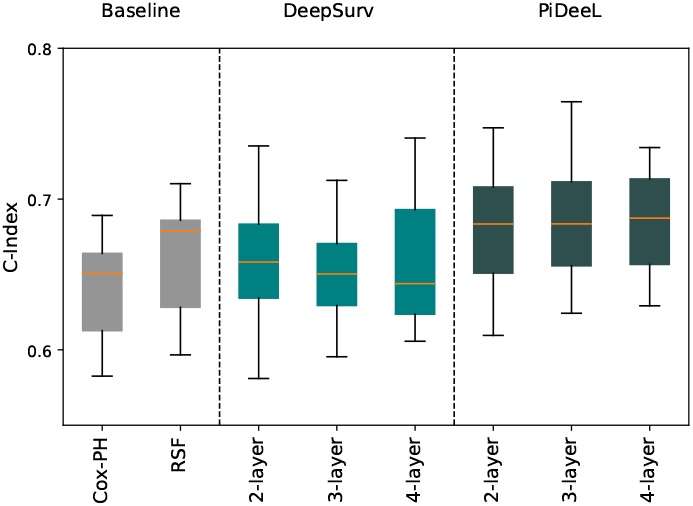
Comparison of baseline models and PiDeeL on survival analysis in terms of c-index. This evaluation is based on 5-fold cross-validation that is repeated 3 times. (i.e., 15 iterations).

### Interpretability and Importance Analyses

When combined with downstream feature importance analysis methods, (SHAP [35]), the pathway-informed architecture of PiDeeL enables us to jointly analyze the importance of metabolites and pathways for the model output. In addition to the quantitative analyses presented above, we perform feature importance analyses on the 4-layer PiDeeL to empirically pinpoint metabolites and pathways candidate for being implicated in gliomas, their progression, and associated patient survival. We calculate SHAP values on the input features (i.e. in silico predicted metabolite concentrations – 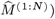 as well as pathway-informed layer activations (*PA*^(1:*N*)^).

Please see Supplementary Figures S4 and S5 where we show PiDeeL’s metabolite importance plots for the survival prediction and tumor pathological classification tasks. We highlight the top 3 metabolites having the highest mean SHAP value for the survival analysis among all 37 metabolites according to our analyses in the decreasing order of importance *alanine, glutamine*, and *glutamate*. We also pinpoint the top 3 pathways having the highest SHAP values among the 138 pathways used for pathway-informed layer, in the decreasing order of importance as: *Mineral absorption, mTOR signaling pathway* and *Alanine - aspartate and glutamate metabolism*.

Through a case-control study, alanine was reported to be a potential biomarker for malignant gliomas [20]. Similarly, glutamate was also found to play a central role in malignant gliomas through multiple mechanisms [14]. Since malignant gliomas usually have a poor prognosis and survival, the high ranking of alanine and glutamate in the list is expected. Furthermore, glutamine is the primary precursor of glutamate and plays a critical role in brain function [37]. Glutamine metabolism can regulate multiple pathways including energy production and redox homeostasis in brain cancer cells [37]. Glutamine itself was pinpointed as a candidate biomarker for glioma progression and response to treatment [16]. Thus, the rank of glutamine in the list is also expected.

We biologically motivate the connection between the top 3 pathways and gliomas. Minerals and their absorption are fundamentally important to sustain life [1]. For example, calcium plays a wide variety of roles in the human body. Moreover, various natural dietary components were reported to have potential benefits in the prevention and treatment of gliomas [31]. Calcium was reported to be one of the dietary components having a potential for glioma prevention by affecting apoptosis and DNA repair [50]. Thus, the rank of *Mineral absorption* pathway according to calculated importance is biologically grounded. Furthermore, the mammalian target of rapamycin (mTOR), is reported to have a key role in integrating signal transduction and metabolic pathways in glioblastoma [2]. Since *∼* 46% of the cohort analyzed in this work were diagnosed with glioblastoma, the importance rank of *mTOR signaling pathway* is viable. Finally, two of the three metabolites implicated in *Alanine— aspartate and glutamate metabolism* pathway, were also detected among the top 3 important metabolites with respect to metabolite-level importance scores, as discussed above. This qualitatively reinforces the predictive capabilities of PiDeeL and showcases its interpretability in both metabolite and pathway levels.

### Ablation Studies for Pathway-Informed Architecture

In this section, we present the results of ablation studies discussed in the Ablation Studies for Pathway-Informed Architecture under the Methods Section. Figure 3 demonstrates the result of the respective itemized ablation test (excluding dropout).

**Fig. 3.**
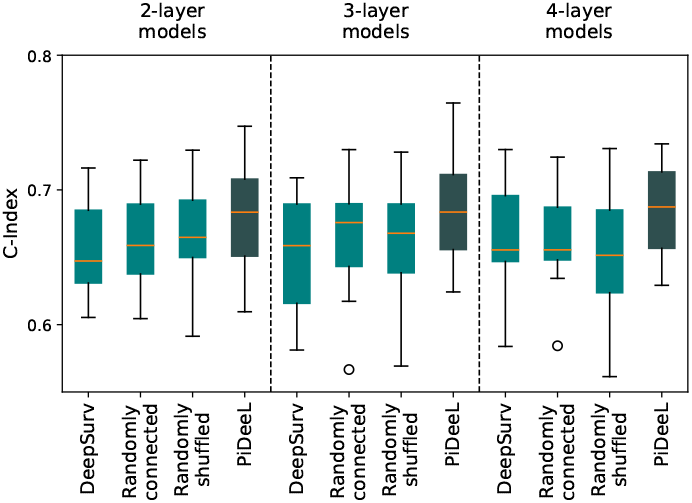
Performance comparison of PiDeeL on survival analysis task against DeepSurv with 138 neurons in the first hidden layer, PiDeeL with randomly connected *PI_matrix*, and PiDeeL with shuffled *PI_matrix*, with respect to c-index.

We divide Figure 3 into 3 panels to show the results of 2, 3, and 4-layer models. Under each panel, the boxplots from left to right demonstrate the performance attained by 3 ablation tests (cyan boxplots) that are DeepSurv model with 138 neurons in the first layer, PiDeeL with randomly connected *PI_matrix*, PiDeeL with randomly shuffled *PI_matrix* and also PiDeeL (navy blue boxplots).

We observe that PiDeeL improves the median c-index over DeepSurv with 138 neurons in the first layer by 3.6%, 2.6%, and 3.6% for 2, 3, and 4-layered networks, respectively. We also see that using PiDeeL over DeepSurv decreases the variance by an average of 29.7%. Note that the original DeepSurv model has 64 neurons in the first layer. This shows that not the larger number of neurons in the first layer but the pathway information leads to the performance improvement over DeepSurv.

We compare the performance of PiDeeL with *PI matrix* against PiDeeL with random *PI_matrix*. In this setup, we randomly connect our metabolite nodes (input vector) and pathway activation layer neurons. For this, we constructed randomly connected 37×138 matrices preserving the total number of connections in the original *PI_matrix* (468). We observe that PiDeeL improves the median c-index over PiDeeL with random *PI matrix* by 3.7%, 1.2%, and 4.9% for 2, 3, and 4-layered networks, respectively.

We show the performance comparison of PiDeeL with *PI matrix* against PiDeeL with shuffled *PI matrix*. This time, instead of randomly connecting the *PI_matrix*, we shuffle the rows to preserve each metabolite-to-pathway profile. We observe a similar trend as in Panel C. This time, *PI_matrix* improves the performance attained by using shuffled *PI_matrix* in terms of median c-index by 2.8%, 2.4%, and 4.9% for 2, 3, and 4-layered networks, respectively. Additionally, using *PI_matrix* decreases the variance by an average of 4.6%.

Figure 4 shows the survival prediction performance comparison of DeepSurv with various dropout rates and PiDeeL. We observe that PiDeeL substantially outperforms DeepSurv-with-dropout regardless of the rate. We also see that DeepSurv’s performance deteriorates when the dropout rate is increased. The 2-layer PiDeeL model achieves a median c-index of 68.3%, in contrast, the DeepSurv model attains median c-index scores of 58.1%, 58.6%, 51.6%, 51.1% and 47.0% for 0.5, 0.6, 0.7, 0.8 and 0.9 dropout rates, respectively. We observe the same trend as we increase the number of layers for both models. This indicates that sparsity provided by the dropout does not contribute to the performance of the fully-connected model for survival prediction, whereas PiDeeL’s pathway-informed sparsity with *PI matrix* improves the performance.

**Fig. 4.**
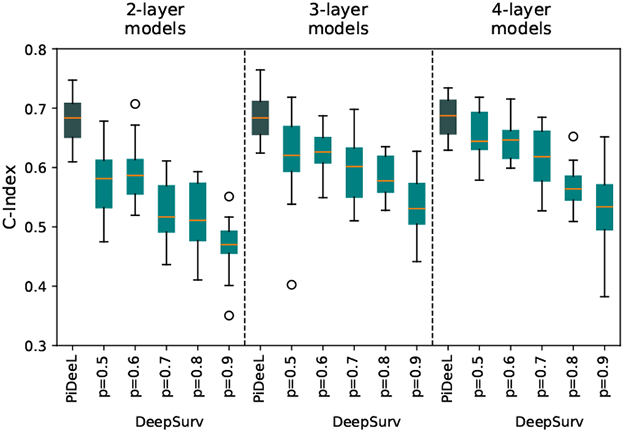
Performance comparison of PiDeeL on survival analysis task against DeepSurv with different dropout rates with respect to c-index. This evaluation is based on 5-fold cross-validation that is repeated 3 times. (i.e., 15 iterations).

## Discussion

1H HRMAS NMR spectrum has been a reliable resource for distinguishing malign and healthy samples taken from the excision cavity during surgery. It is a good fit because it is non-destructive and can analyze small samples of raw tissue specimens [22]. Machine learning techniques that learn from the NMR signal to distinguish healthy and tumor tissue, as well as benign and aggressive tumors, have been an important tool. Yet, the performance was limited due to small training set sizes [7, 8] which prohibit using complex architectures like deep neural networks learning a hierarchical composition of complex features. In this study, we are able to use such a hierarchically complex model for distinguishing benign and aggressive tumors for the first time with comparable performance. Despite having a deep model, we are able to decrease the number of trainable parameters in the model by composing the architecture with prior biological information obtained from metabolic networks.

The pathology of the tumor is an important cue for the surgeon who might want to resect more tumor tissue and risk the patient’s well-being if the tumor is likely aggressive, or who might risk leaving some residual tumor tissue if the tumor is likely benign. Yet, this does not one-to-one determine the prognosis. Thus, it is also important to estimate the survival of a patient, which is a proxy for prognosis, and inform the surgeon. For this, we train a pathway-informed model to predict the survival of patients and show that we attain high performance. An example showing the usefulness of the survival prediction along with pathological classification is as follows: Our model is able to correctly predict long survival for a patient with aggressive glioma, who lived 5,656 days after surgery. The surgeon can be more conservative in this case compared to being informed with only a prediction of an aggressive tumor. Similarly, our model correctly predicts low survival for a patient with a benign tumor who lived only 43 days after surgery.

We also investigate the use of a multi-task learning architecture that combines two architectures and predicts both labels simultaneously. Our results demonstrate that this approach only slightly increases the performance for both tasks. For this reason, we ultimately decided to proceed with the utilization of two task-specific single-task PiDeeL models.

One limitation of our study is that it is bottlenecked by the automated metabolite quantification model which has been shown to have promising performance and generalizability. Yet, this component consists of individual multi-layered perceptrons for 37 metabolites for which a training dataset exists. We think our model would benefit from a more comprehensive spectrum of metabolites to predict the survival of a patient which is a complex outcome to approximate. The human metabolome roughly consists of 3,000 metabolites and we are currently limited to using only 1% of this information source.

## Competing interests

No competing interest is declared.

## Acknowledgments

This work was supported by grants from BPI France (ExtempoRMN Project), Hôpitaux Universitaires de Strasbourg, Bruker BioSpin, Univ. de Strasbourg, and the Centre National de la Recherche Scientifique; also by TUBA GEBIP, Bilim Akademisi BAGEP and TUSEB Research Incentive awards to AEC.

## Notes

### Competing Interest Statement

The authors have declared no competing interest.

### Summary of Updates

We have revised the ablation study. We corrected various grammar mistakes and revised the text for better readability.

