## Supplementart Material for "PiDeeL: Pathway-Informed Deep Learning Model for Survival Analysis and Pathological Classification of Gliomas"

#### 1 Supplementary Text

##### 1.1 Dataset

###### 1.1.1 Tissue Sample Collection

All tissue specimens were collected either by a pneumatic system connecting the operating theater of neurosurgery to the NMR room (Hautepierre hospital, University Hospitals of Strasbourg) or from samples stored at Strasbourg and Colmar tumor biobanks, with minimum ischemic delays (average time $2\,\mathrm{min} \pm 1\,\mathrm{min}$). The collected samples were adequate for proper HRMAS NMR analysis and had a low necrosis-to-tumor ratio. After HRMAS NMR analysis, inserts for half the content of each sample were cut. The percentage of tumor cells in the sample was calculated with respect to the total surface area as well as frozen hematoxylin & eosin-stained sections. Please see Supplementary Table 1 for the histopathological classification of the tumor tissue samples.

###### 1.1.2 Ethics Statement

The tissue specimens used in this study were collected from samples stored in Strasbourg and Colmar tumor biobanks, or by a pneumatic system. Collection, storage, and compilation of the dataset were approved by the Ethics Committee no. 2003-100., Ethics Committee no. 2013-37, 12.11.2013. and Ethics Committee of Strasbourg, (Comité de protection des personnes, Est IV), respectively. Consent for all patients who participated in this study was acquired by a written letter.

###### 1.1.3 HRMAS NMR Spectrum Acquisition

A Bruker Avance III 500 spectrometer equipped with a 4-mm triple-resonance gradient HRMAS probe (1H, 13C, and 31P) operating at a proton frequency of 500.13 MHz was used to acquire the spectra from biopsy samples prepared by combining $30\mu L$ KelF insert and 15- to 18-mg collected sample at 20°C. The lock frequency of the spectrometer was set by adding $10\mu L$ of $D_2O$ to the insert. Spectrum acquisition was performed using a Carr-Purcell-Meiboom-Gill (CPMG) pulse sequence with a $285\mu s$ inter-pulse delay and 10 min acquisition time. The CPMG pulse train had a length of 93 ms due to 328 loops. During the spectrum acquisition process, the temperature was stabilized to 4°C in order to reduce the effects of tissue degradation. As a result, a one-dimensional (1D) proton (1H) spectrum was acquired for each specimen. Finally, a simulated,

Lorentzian-shaped ERETIC [?] signal, whose amplitude was calibrated to correspond to exactly 69.6 nmoles of protons, was added to the spectrum digitally at 10 ppm. The amplitude of the digitally added ERETIC signal also takes into account the difference in pulse widths between the standard acetate sample and the biopsy sample under investigation, making the method insensitive to the salt effect [?]. The calibrated ERETIC signal allowed the quantification of all the identified metabolites present in the spectrum.

#### 1.2 Automated Metabolite Quantification

The automated metabolite quantification pipeline method presented in [?] was used to quantify the following 37 metabolites of interest: 2-hydroxyglutarate, 3-hydroxybutyrate, Acetate, Alanine, Allo-cystathionine, Arginine, Ascorbate, Aspartate, Betaine, Choline, Creatine, Ethanolamine, GABA, Glutamate, Glutamine, Glutathione (GSH), Glycerophosphocholine, Glycine, hypo-Taurine, Isoleucine, Lactate, Leucine, Lysine, Methionine, Myo-inositol, NAL, NAA, o-Acetylcholine, Ornithine, Phosphocholine, Phosphocreatine, Proline, scyllo-Inositol, Serine, Taurine, Threonine and Valine.

In order to prepare our HRMAS NMR spectra for this pipeline, we first remove the 70 initial time points in the FID signal which correspond to the spectroscopy machine-specific (Bruker for our case) digital filter. The clipped time domain signal was transformed to the frequency domain via Discrete Fourier Transform without windowing and phase-corrected after a line broadening of 10Hz was applied. The magnitude of the resulting signal (ranging from -2ppm to 12ppm) was normalized by tissue sample mass and multiplied with the constant acetate reference-based scaling factor provided in their paper. Finally, the scaled magnitude signal was binned to $1,401$ bins of 0.01 ppm-width by summing the intensity of the frequencies mapped to each bin. The resulting signal was used as input to the publicly available trained 2-layer fully-connected neural network-based quantification models available for each metabolite of interest from `https://github.com/gunkaynar/targeted_brain_tumor_margin_assessment`.

#### 1.3 Experimental Setup

##### 1.3.1 Hyperparameter Space

In this section, we give the lists of hyperparameter space for baseline models for both survival analysis and pathological classification tasks. In the of CoxPH model, we use a ridge-regularized variant to prevent overfitting. We perform hyperparameter optimization on the validation split. The hyperparameter search space was limited to: (1) regularization penalty coefficient $\in \{0, 0.1, ..., 0.9, 1\}$; (2) the heuristic for breaking ties in event ties was set to *efron* [?] or *breslow* [?]; and (2) the maximum number of iterations $\in \{100, 200, ..., 500\}$.

For the RSF model, similar to the case with Cox-PH, we perform hyperparameter selection on validation split using the following search space: the number of trees in the forest $\in \{20, 50, 100, 150, 300, 400]\}$; the maximum depth limit for each tree $\in \{no\ limit, 10, 15, 25, 30\}$; the minimum number of samples required to split an internal node in a tree $\in \{2, 5, 10, 15\}$; the minimum number of samples required to be available in leaf nodes resulting from considered split $\in \{2, 5, 10, 15\}$; and the maximum number of features explored while looking for best splitting criterion $\in \{sqrt(number\ of\ metabolites = 37), log_2(37)\}$.

DeepSurv model's hyperparameter search space is limited to: (1) batch size for training $\in \{16, 32, 64, 128, full - batch\}$; and (2) activation function between layers $\in \{ReLU, Sigmoid\}$.

On the other hand, for the pathological classification task, we have RF as the baseline model. In the case of RF models, we apply grid search for the hyperparameter selection and selected the setting with the best-observed performance (AUC-ROC) on the validation set. Hyperparameter search space for the RF model is as follows: the number of trees in the forest $\in \{50, 150, 300, 400]\}$; the maximum depth limit for each tree $\in \{10, 15, 25, 30\}$; the minimum number of samples required to split an internal node in a tree $\in \{5, 10, 15\}$; the minimum number of samples required to be available in leaf nodes resulting from considered split $\in \{2, 10, 20\}$; criterion to measure the quality of a split: *Gini-index* and *entropy*.

The other baseline model for the pathological classification task is fully-connected neural networks. The hyperparameter search space is similar to DeepSurv as follows: (1) batch size for training $\in \{16, 32, 64, 128, full-batch\}$; and (2) activation function between layers $\in \{ReLU, Sigmoid\}$.

##### 1.3.2 Hyperparameter Selection

Regardless of the task, we use grid search on validation split to determine the best-performing hyperparameter values per model and iteration. Please see section 1.3.1 of Supplementary Text for the hyperparameter space in the case of baseline models. Training these models involved a two-step process. During iteration $i$ of the cross-validation setup, these models were tested on the fold $i$, validated on fold $(i + 1) mod 5$, and trained on the remaining 3 folds. After the selection of the best-performing hyperparameter setting based on the c-index calculated on the validation set, we re-trained the selected configuration on the union of training and validation splits to get stronger baseline models.

Hyperparameters of deep learning-based models explored (DeepSurv) or proposed (PiDeeL) for survival analysis were tuned based on CoxPH loss calculated on validation split predictions. Given its formulation in the Survival Analysis Section, CoxPH loss (Eq. 4) we adopted for deep learning-based survival analysis is a differentiable surrogate for time-dependent c-index metric [?]. Similar to the case with the survival models, hyperparameters of deep learning models for the pathological classification task were also tuned based on binary cross-entropy loss (which is also a differentiable proxy for AUC-ROC and AUC-PR) calculated on validation split. For all deep learning models discussed above, cross-validation folds were distributed to training, validation and test splits similar to CoxPH and RSF models. However, re-training with the union of training and test splits step was not performed due to the presence of early-stopping dynamics. To form the hyperparameter space, we sampled 5 equidistant learning rates from $[10^{-3}, 10^{-4}]$ interval, performed experiments, and proceeded with $10^{-4}$ as the learning rate due to the highest empirically observed performance.

The hyperparameter search space for PiDeeL for both tasks was as follows: (1) batch size for training $\in \{16, 32, 64, 128, full-batch\}$; and (2) activation function between layers $\in \{ReLU, Sigmoid\}$. After conducting thorough experimentation with these hyperparameters, we determined the optimal settings for our model. Specifically, we selected batch sizes of 64 and 32 for survival analysis and pathological classification tasks, respectively. Additionally, we found that the use of the ReLU activation function resulted in the highest empirical performance, and thus, we proceeded with this choice for the activation function.

Moreover, we trained models at most $5,000$ epochs (selected from $1000, 2000, ..., 6000$) and applied early-stopping when validation loss increased for 150 consecutive epochs (selected among 50, 100, 150, 200). We used Adam optimizer [?] with a weight decay of 0.002, also selected based on the highest empirically observed performance.

##### 1.3.3 Implementation Details

The codebase for this work is implemented in Python and is publicly available at `https://github.com/ciceklab/PiDeeL`. Specifically, RSF and CoxPH models were used from [?], and deep learning models were developed with the help of PyTorch framework [?]. All models are trained and tested on a SuperMicro SuperServer 4029GP-TRT with 2 Intel Xeon Gold 6140 Processors (2.3GHz, 24.75M cache), 251GB RAM, 6 NVIDIA GeForce GTX 2080 Ti GPUs (11GB, 352Bit) and 2 NVIDIA TITAN RTX GPUs (24GB, 384Bit). Training of CoxPh, RSF, and DeepSurv models took 1.5, 14, and 2 minutes on average. The training of PiDeeL took 1.5 minutes on average.

#### 1.4 PiDeeL

##### 1.4.1 Pathological Classification

We assess the performance of the pathological classification models with respect to the Area Under the ROC Curve (AUC-ROC) and the Area Under the Precision-Recall Curve (AUC-PR) metrics. Please see Supplementary Figure S2 for a visual depiction of pathological classification performance attained by RF (gray boxplot), fully-connected networks (cyan boxplots), and PiDeeL (navy blue boxplots) with 2, 3, or 4 layers. Here, similar to the survival analysis results presented in the Survival Analysis Section under the Results of the manuscript, we observe that PiDeeL consistently improves upon the fully-connected baseline, regardless of architecture depth or the performance metrics used for the current evaluation. In the case of AUC-ROC, we see 1.7%, 0.7%, and 2.7% improvement in performance PiDeeL compared with the fully-connected baseline with 2, 3, and 4 layers. As implicitly mentioned above, we see a similar trend for the AUC-PR metric where improvements are 0.6%, 0.4%, and 0.8% for the same order of model depth. However, for the current setup of the pathological classification task, we interestingly do not quantitatively observe a big improvement with PiDeeL over the RF model. We observe that when the number of layers in fully-connected networks increases, the performance of the pathological classification deteriorates. However, for PiDeeL, this does not apply to the AUC-ROC metric where the median AUC-ROC slightly increases with the depth of the model.

##### 1.4.2 Multi-task Learning

We explore a multi-task strategy for PiDeeL for survival analysis and pathological classification tasks. The multi-task PiDeeL learns a function $s : \mathbb{R}^{37} \rightarrow [-\infty, \infty] \times \{0, 1\}$. Note that we use the same feature vector (metabolite profiles) as input. The multi-task PiDeeL model architecture can be summarized as follows:

$$PL^{(138)} = \left[\ FC^{(138)}weights\ \right]\left[PI\_matrix\right] \tag{1}$$

$$PA^{(1:N)} = ReLU(PL^{(138)}(\widehat{M}^{(1:N)}) + b^{138}) \tag{2}$$

$$X_{hid}^{(1:N)} = ReLU(FC^{(64)}(PA^{(1:N)})) \tag{3}$$

$$\widehat{h}^{(1:N)}, \widehat{Y_m}^{(1:N)} = FC^{(2)}(X_{hid}^{(1:N)}) \tag{4}$$

We train 2,3 and 4-layer multi-task PiDeeL with the negative log of Cox Partial Likelihood and class-weighted binary cross-entropy losses. Additionally, hyperparameter configurations of multi-task PiDeeL were selected based on optimizing a linear combination of the survival and pathological classification surrogates. Specifically, a grid search based on validation performance led to a 0.125-fold weight for COX-PH loss and a 0.875-fold weight for binary cross-entropy loss. Please see Supplementary Figure S6 for a visualization of the convergence of the loss functions during training for a 3-layer PiDeeL model using a random seed. The left plot shows the absolute values of the Cox-PH and binary cross-entropy losses, while the right plot shows the scaled values of Cox-PH and binary cross-entropy losses.

Please see Supplementary Figure S3 for the performance comparison of single and multi-task learning of survival analysis and pathological classification tasks using PiDeeL. Across all analyzed model depths, we observe that the multi-task modeling strategy is at least slightly beneficial for performance on the survival analysis task. The 4-layer multitask PiDeeL attains the highest median c-index of 69.5%. We observe that the 2-layer multi-task PiDeeL outperforms all single-task and multi-task models in the pathological classification task with a median AUC-ROC of 91.9% and a median AUC-PR of 97.5%. Nonetheless, the examination of multi-task learning of PiDeeL yielded a slight improvement in performance (0.8% in median c-index and 1.8% in median AUC-ROC and 0.4% in median AUC-PR, in 4-layer and 2-layer multi-task PiDeeL models); thus, we decide to proceed with the utilization of two task-specific single-task PiDeeL models.

##### 1.4.3 Interpretability and Importance Analyses

In this section, we cover the details of interpretability and importance analyses performed on PiDeeL for metabolite and pathway-activation vectors. First, we adopt the same 5-fold cross-validation fold setup (repeated 3 times) and hyperparameter selection described in the Experimental Setup Section of the manuscript. During iteration $i$, SHAP values are calculated on the test fold (fold$(i+1)mod5$) on the trained PiDeeL model of training fold $i$. The hyperparameters are preserved among all trained models (5-fold repeated 3 times). In each iteration (i.e., 15 times), we concatenate the SHAP values calculated in this iteration with the previous SHAP values to get a $37x15$ matrix. We averaged each metabolite's SHAP value vector and sort the 37 metabolite's mean SHAP values in increasing order. Please see Supplementary Figures S4 and S5 for the calculated SHAP values for each metabolite for survival analysis and pathological classification tasks, respectively. From this analysis, 3 metabolites with the biggest impact on PiDeeL's prediction on survival analysis are discussed in Section 3.3 of the manuscript.

After that, to assess the interpretability of the pathway profiles, we use the same 5-fold cross-validation setup. But this time, we take the first hidden layer's output (i.e., pathway-activation vector $PA^{(1:N)}$). We measure the impact of each pathway by calculating the SHAP values of vector $PA^{(1:N)}$ on PiDeeL's prediction. We again concatenate and sort SHAP values of $PA^{(1:N)}$) calculated on each fold. The 3 most important pathways are discussed in the Interpretability and Importance Analyses Section of the manuscript.

##### 1.4.4 Ablation Studies

In this section, we give the details of the ablation tests we performed to identify the contribution of the pathway-informed architecture to the performance of the PiDeeL model.

In test (a), we validate the contribution of the number of neurons in the first layer to the performance. Note that in PiDeeL, the first hidden layer has 138 neurons, whereas trained fully-connected networks (baseline models) have 64. We train 2, 3, and 4-layer DeepSurv with the same 5-fold cross-validation setup, and repeat it 3 times. The model architecture can be summarized as follows:

$$X_{hid}^{(1:N)} = ReLU(FC^{(138)}(\widehat{M}^{(1:N)})) \tag{5}$$
$$X_{hid}^{(1:N)} = ReLU(FC^{(64)}(X_{hid}^{(1:N)})) \tag{6}$$
$$\widehat{h}^{(1:N)} = FC^{(1)}(X_{hid}^{(1:N)}) \tag{7}$$

Second, in test (b), we validate the contribution of the first hidden layer's sparsity to the performance of PiDeeL. The total connections between the input layer ($\widehat{M}^{(1:N)}$) and the first hidden layer $PL^{(1:N)}$ are 468 in PiDeeL, whereas in a fully-connected network with 37 input features, there are 5106 (considering the case of 138 first hidden layer neurons). We train 2, 3, and 4-layer DeepSurv with dropout in the first hidden layer adopting the same 5-fold cross-validation setup, and repeat it 3 times. We train fully-connected networks using different dropout rates from $\{0.5, 0.6, 0.7, 0.8, 0.9\}$.

Recall that $PL^{(138)}$ refers to the weights of the pathway-informed layer of PiDeeL. To construct the $PL^{(138)}$, we use Eq. 8 where we multiply $FC^{(138)}weights$ with $PI\_matrix$. Note that we build $PI\_matrix$ from the KEGG database for pathways. In test (c), we use randomly connected $PI\_matrix$ (to verify the contribution of $PI\_matrix$ to the performance of PiDeeL. The number of connections in the first hidden layer (468) is preserved in random $PI\_matrix$.

$$PL^{(138)} = \left[\ FC^{(138)}weights\ \right]\left[PI\_matrix\right] \tag{8}$$

We train 2, 3, and 4-layer PiDeeL with the same 5-fold cross-validation setup, and repeat it 30 times (i.e., 150 iterations).

Finally, in test (d), we construct $PI\_matrix$ similar to test (c). However, we shuffle each metabolite-to-pathway connection vector instead of randomly connecting. Note that $PI\_matrix$ is a $37x138$ matrix; and for metabolite $j$, $PI\_matrix.j$ yields a vector of shape $1x138$. We shuffle these 37 vectors and train 2, 3, and 4-layer PiDeeL with the same 5-fold cross-validation setup, and repeat it 30 times (i.e., 150 iterations).

Please see Figure 3 in the manuscript for the depiction of the results of ablation studies. Please refer to the Ablation Studies for Pathway-Informed Architecture Section of the manuscript for the observations of ablation tests for the survival analysis task.

#### 2 Supplementary Figures

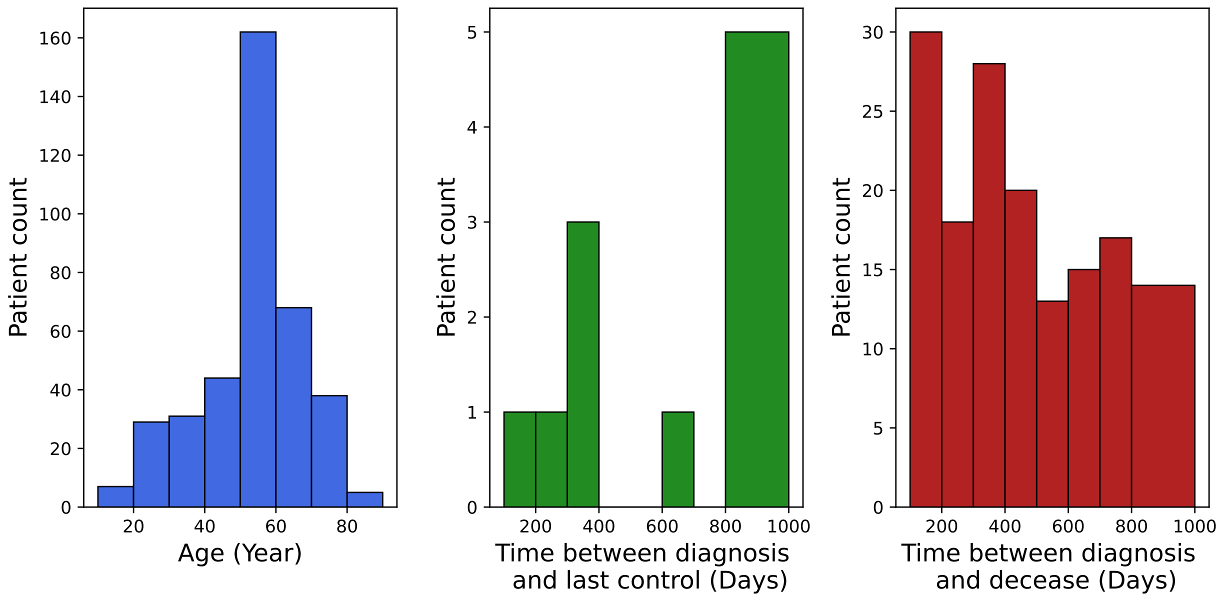


Supplementary Figure 1: (i) the age distribution of patients in the dataset, (ii) the distribution of duration between the primary tumor removal surgery and the last control, (iii) the duration distribution of duration between the primary tumor removal surgery and decease

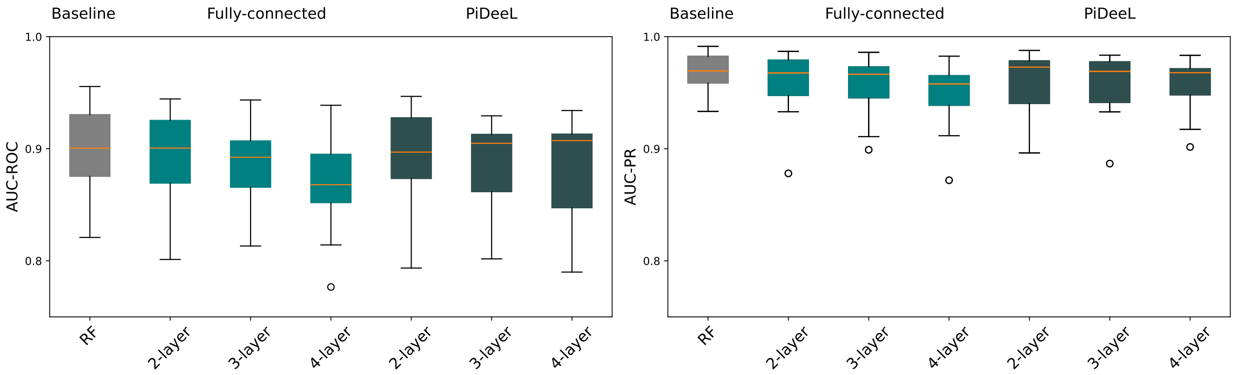


Supplementary Figure 2: Comparison of baseline models and PiDeeL on pathologic classification in terms of AUC-ROC (left) and AUC-PR (right). This evaluation is based on 5-fold cross-validation that is repeated 3 times. (i.e., 15 iterations)

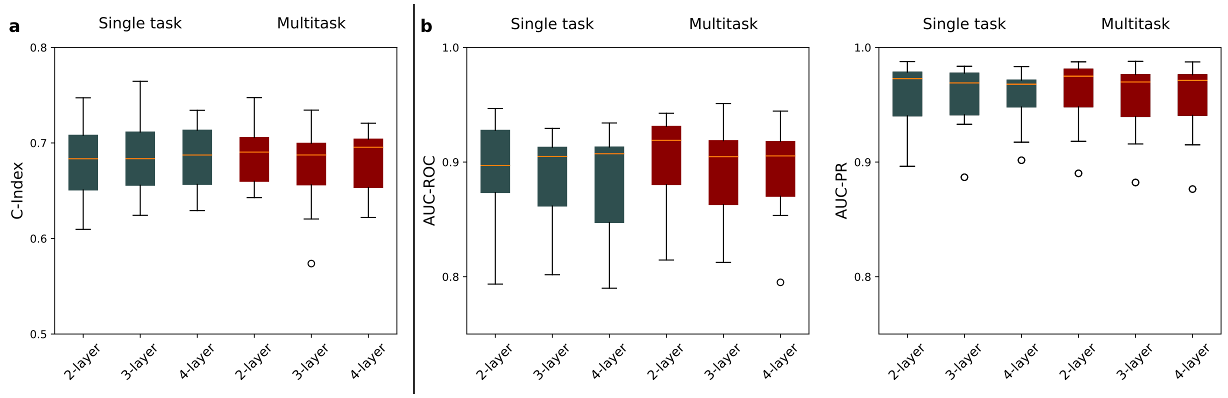


Supplementary Figure 3: The performance comparison between task-specific PiDeeL (navy blue boxplots) versus multi-task learning of PiDeeL (red boxplots). The survival analysis performance is under Panel A and the pathological classification performance is under Panel B, with AUC-ROC on the left, and AUC-PR on the right. This evaluation is based on 5-fold cross-validation that is repeated 3 times. (i.e., 15 iterations)

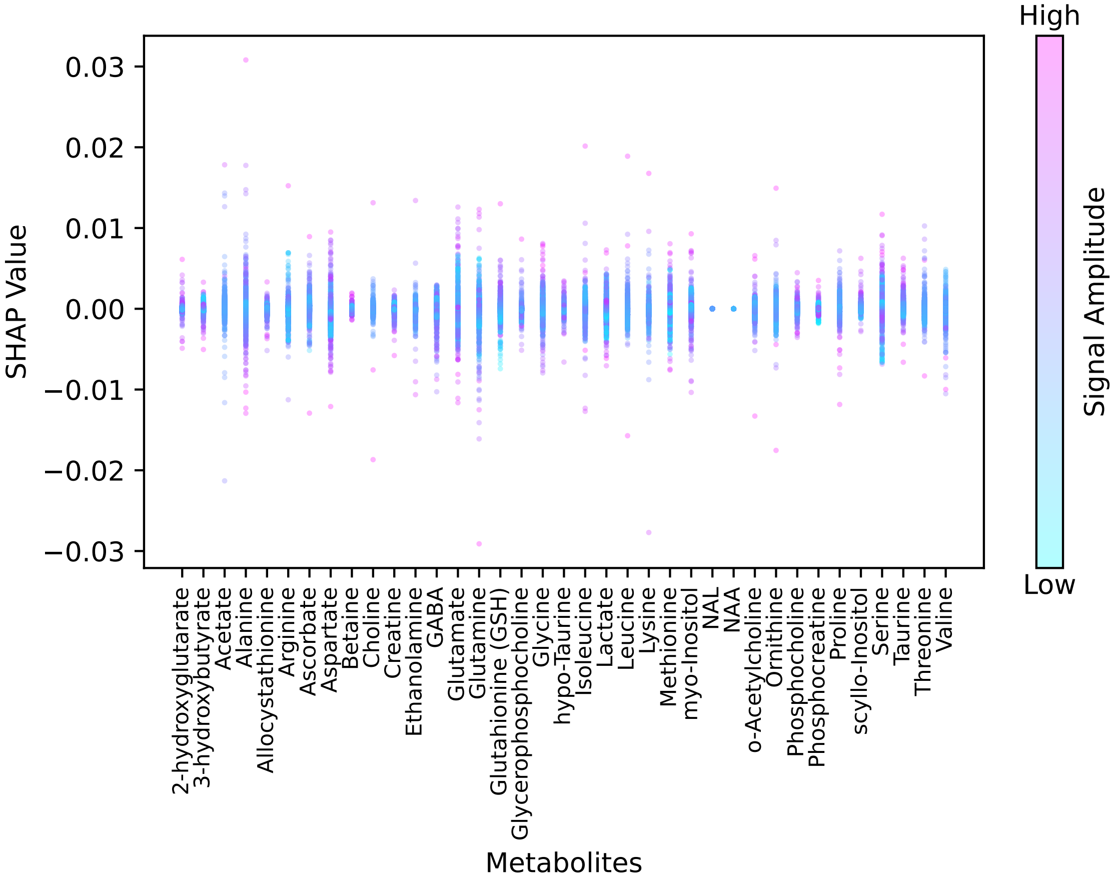


Supplementary Figure 4: The SHAP values (on the y-axis) for the survival analysis task of each quantified biomarker metabolite (on the x-axis) are shown. On the left, the color bar is shown (purple represents higher value and blue represents lower value for feature). The more the SHAP value, the more critical the metabolite is for the model's output toward a higher risk score for survival analysis.

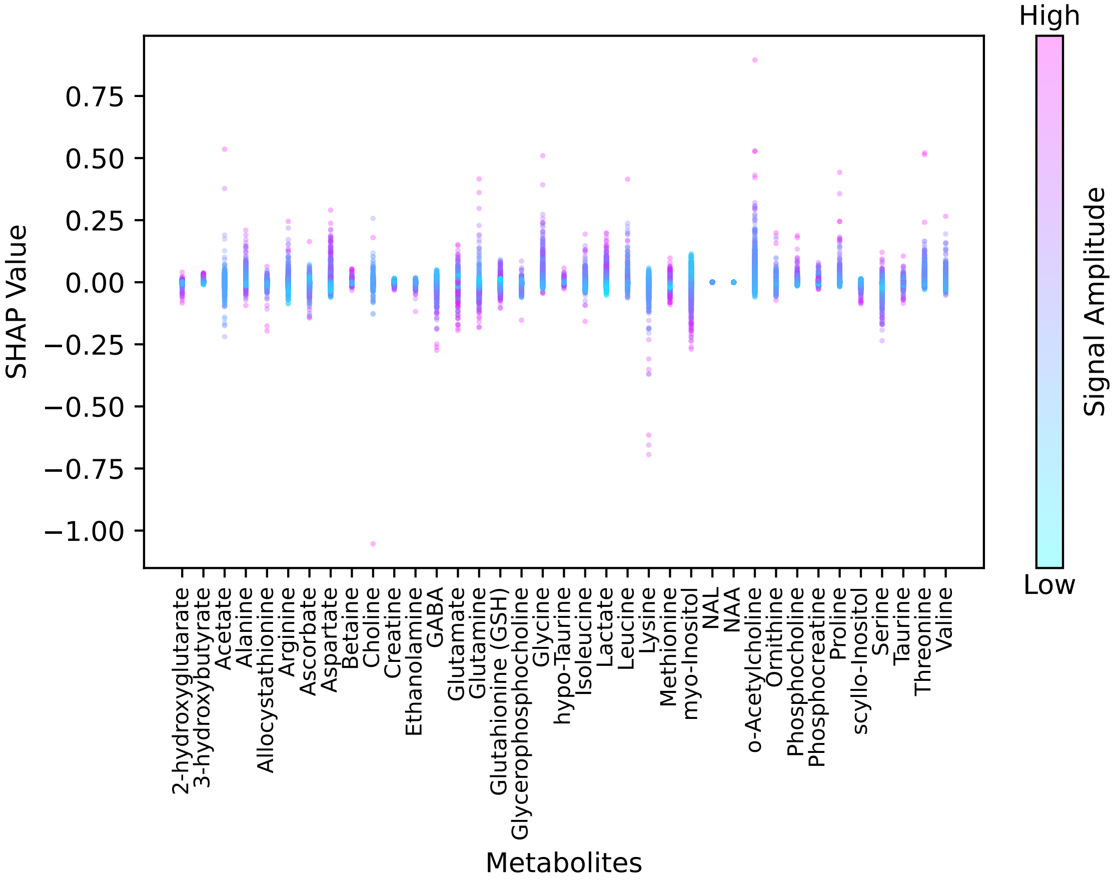


Supplementary Figure 5: The SHAP values (on the y-axis) for the pathological classification task of each quantified biomarker metabolite (on the x-axis) are shown. On the left, the color bar is shown (purple represents higher value and blue represents lower value for feature). The more the SHAP value, the more critical the metabolite is for the model's output toward a higher risk score for survival analysis.

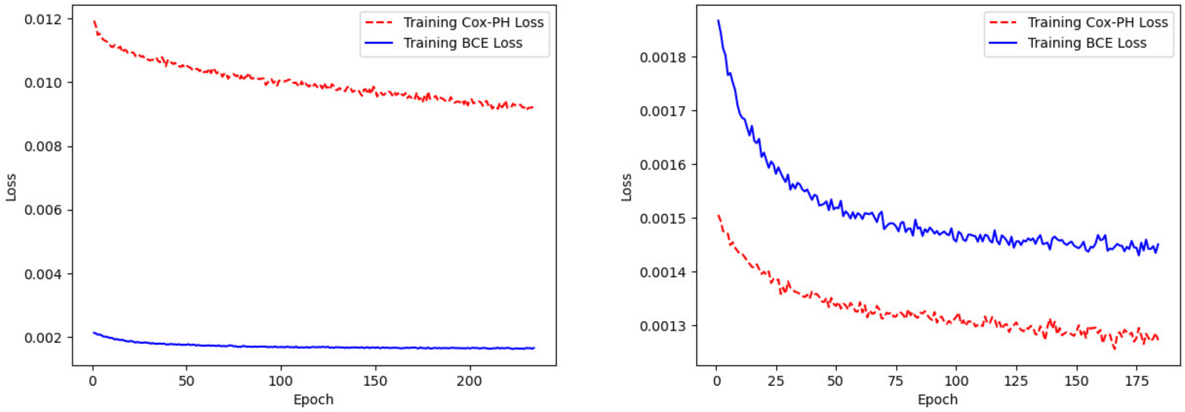


Supplementary Figure 6: The training loss convergence during training for the 3-layer PiDeeL model using a random seed. On the left, the absolute values of Cox-PH and BCE losses are shown, whereas, on the right, the scaled loss values are shown. The weights for Cox-PH and BCE losses are 0.125 and 0.875, respectively.

### 3 Supplementary Tables

| **Primary Brain Tumor Types** | **Number of Samples** |
|---|---|
| Pilocytic astrocytoma (AST-I) | 4 |
| Astrocytoma grade II (AST-II) | 5 |
| Astrocytoma grade III (AST-III) | 7 |
| Glioblastoma (GBM) | 178 |
| Oligodendroglioma grade II (ODG-II) | 36 |
| Oligodendroglioma grade III[1] (ODG-III) | 103 |
| Oligoastrocytoma grade II (OAST-II) | 3 |
| Oligoastrocytoma grade III (OAST-III) | 9 |
| Ganglioma grade II (GG-II) | 5 |
| Ganglioma grade III (GG-III) | 1 |
| Dysembryoplastic Neuroepithelial Tumors (DNET) | 25 |
| Gliosarcoma (GS) | 3 |
| Unkown subtype | 5 |

Supplementary Table 1: Histopathological classification of the tumor tissue specimens used in the study.

[1]Three samples had glioblastoma as well as oligodendroglioma grade III. They are included in the sample count for the latter.
